# Multiplexed Data-Independent Acquisition (mDIA) to Profile Extracellular Vesicle Proteomes

**DOI:** 10.1101/2025.05.12.653483

**Authors:** Yi-Kai Liu, Nathaniel Miller, Marco Hadisurya, Zheng Zhang, W. Andy Tao

## Abstract

Extracellular vesicles (EVs) have gained increasing attention with their intriguing biological functions and their molecular cargoes serving as potential biomarkers for various diseases, including cancers. A relatively lower abundance of EV proteins compared to cellular counterparts necessitates sensitive and accurate quantitative proteomic strategies. Multiplexed proteomics combined with data-independent acquisition (mDIA) has shown promise for improving sensitivity and quantification over traditional DDA and label-free methods. Despite this, mDIA pipelines that utilize various types of spectral libraries and search software suites have not been thoroughly evaluated with EV proteome samples. In this study, we aim to establish a robust mDIA pipeline based on dimethyl labeling for quantitative proteomics of EVs. EVs were isolated using the extracellular vesicle total recovery and purification (EVtrap) technique and processed directly through an on-bead one-pot sample preparation workflow to obtain digested peptides. We evaluated different mDIA pipelines, including library-free and library-based DIA on the timsTOF HT platform. Results showed that library-based DIA, with project-specific spectral libraries generated from StageTip-based fractionation, outperformed other pipelines in protein identification and quantification. We demonstrated for the first time EV proteome landscape changes caused by the IDH1 mutation and inhibitor treatment in intrahepatic cholangiocarcinoma, highlighting the utility of mDIA in EV-based biomarker discovery.

## Introduction

Extracellular vesicles (EVs) have been reported as valuable resources for biomarker discovery due to the molecular cargo they carry, which reflects the physiological or pathological state of their originating cells (1, 2). Studying these cargoes, such as proteins, lipids, nucleic acids, and metabolites, is essential for understanding the role of EVs in cell-to-cell communication in the context of various diseases (3–5). Moreover, EVs can be found in almost all body fluids, including blood, urine, saliva, and cerebrospinal fluid, to transport a variety of biomolecules (6–9). These molecules can signal cellular changes and potentially reveal early indicators of diseases, making EVs a promising source for non-invasive diagnostics (10–12). This potential is particularly significant in diseases such as cancer, where early detection and monitoring of therapeutic response are critical (13, 14). EV proteomics, which focuses on detecting protein profiles within or associated with EVs, has thus gained substantial attention in the research community for its potential in understanding EV biology, biomarker discovery, and disease diagnosis (15, 16).

Despite their promise, EVs present several analytical challenges, primarily due to the relatively low abundance of EV-derived proteins compared to proteins in cells or free-circulating proteins (17–19). This challenge makes it difficult for conventional proteomic methods, which often rely on analyzing more abundant proteins, to detect EV proteins with the necessary sensitivity and representativeness. Furthermore, quantification of the levels of EV proteins requires an efficient and reproducible EV isolation method in order to identify reliable biological relevant changes (18, 20). These issues have spurred the development of more sensitive and accurate quantitative proteomic approaches capable of analyzing low-abundance proteins within EVs.

Data-independent acquisition (DIA) has become a more popular mass spectrometry (MS) technique that offers several advantages over conventional data-dependent acquisition (DDA) (21, 22). Unlike DDA, which focuses on the most abundant ions in a sample, DIA acquires comprehensive data by fragmenting all ions within a predefined mass-to-charge or ion mobility range, enabling more complete proteome coverage. This is particularly advantageous for low-abundance proteins found in EVs, as DIA increases the likelihood of detecting their fragment ions even when they are present at trace levels (23, 24). To further improve quantification outcomes, especially for these low-abundance proteins, sample multiplexing is commonly employed. By allowing multiple samples to be analyzed simultaneously, multiplexing reduces technical variability and facilitates more confident comparisons across experimental conditions, improving the reliability of low-abundance protein quantification (25). This is typically achieved through chemical labeling (e.g., tandem mass tag (TMT), dimethylation) or metabolic labeling (e.g., stable isotope labeling by amino acid in cell culture (SILAC)) of proteins or peptides (26–28). By pooling the samples, a single MS run can distinguish proteins with different isotope labels, allowing for their quantification across various samples. This strategy also enhances the throughput of LC-MS/MS analysis, resulting in more precise and efficient quantification.

Recently, some research groups have explored the potential usage of combining DIA with multiplexing, multiplexed DIA (mDIA), to further enhance detection outcomes (29, 30). Although the nature of mDIA results in highly complex MS spectra, several studies have shown its feasibility by using modern high-resolution MS and advanced search algorithms. For instance, pulsed SILAC has been integrated with DIA analysis to enhance sensitivity and quantification in protein degradation studies (30, 31). Derks et al. combined amine-reactive non-isobaric isotopologous mass tags (mTRAQ) with a DIA framework, achieving exceptional detection sensitivity without requiring offline fractionation for multiplexed samples (32). More recently, Thielert et al. utilized dimethyl labeling-based mDIA in conjunction with the RefQuant algorithm to enable highly sensitive and high-throughput single-cell spatial proteomics (33). With the advantageous features, mDIA can potentially benefit EV proteomic analysis.

However, the use of different types of spectral libraries and search software suites may introduce variability in data analysis, potentially impacting mDIA outcomes (34). Spectral libraries, which contain information on the fragmentation patterns of peptides, are essential for interpreting DIA data. Various types of libraries, such as project-specific and predicted libraries, can be applied to DIA proteomics, each with different strengths and limitations (35, 36). Additionally, different search software suites, each with distinct algorithms and workflows, further complicate the establishment of a standardized mDIA pipeline for EV analysis (37). Since mDIA will generate more complex spectra than label-free DIA, a comprehensive evaluation of different mDIA pipelines is needed to ensure reliable identification and quantification of EV proteins, especially for detecting biologically relevant changes.

In this study, we aim at establishing a robust mDIA pipeline for quantitative EV proteomics. EVs were isolated through the EVtrap technique and fully characterized according to the Minimal Information for Studies of Extracellular Vesicles (MISEV) guidelines (38–40). Several data acquisition pipelines, including conventional DDA, library-free DIA, and library-based DIA, were compared on the timsTOF HT platform. Our results indicated that library-based DIA, with project-specific spectral libraries generated from StageTip-based fractionation, provided superior protein identification and quantification. Using the optimized pipeline, we detected proteome changes associated with isocitrate dehydrogenase 1 (IDH1) mutations in intrahepatic cholangiocarcinoma (ICC) cell-derived EVs. The pipeline also identified proteins in EVs from IDH1 mutant cancer cells responding to IDH1 inhibitor AG-120 treatment. Overall, our study established an effective mDIA pipeline for sensitive and accurate EV proteomics and demonstrated, for the first time, the effect of IDH1 mutation and drug treatment on ICC EV proteome, underscoring the potential of mDIA in EV biomarker discovery.

## Experimental Procedures

### Cell culture and media collection

Two ICC cell lines, CCLP (IDH1 wildtype) and RBE (IDH1 mutant), were cultured in RPMI-1640 medium supplemented with 10% heat-inactivated fetal bovine serum, 100 μg/mL streptomycin, and 100 IU/mL penicillin in a 5% CO₂ at 37°C. Once cells reached over 90% confluency, they were switched to serum-free media following three washes. For the inhibitor treatment condition, RBE cells were treated with 100 nM AG-120 for 5 days (41), with a single media change on day 3. The cells were then switched to serum-free media. After 24 hours, the media were collected into 15 mL conical tubes, and cells were washed three times with 1× PBS before collection. All media were then centrifuged separately at 300 × g for 5 minutes to remove floating cells, followed by 2,500 × g for 10 minutes to eliminate debris, large particles, and aggregates. The cell pellets and media supernatants were frozen and stored at -80°C for future use.

### EV isolation and peptide sample preparation

The standard EVtrap protocol was performed according to the manufacturer’s instructions. For each sample, 15 mL of media were used for EV isolation. The isolated EVs on the beads were subjected to a one-pot sample preparation procedure according to the previous study with some modifications (42). In brief, 50 μL of lysis solution containing 12 mM sodium deoxycholate, 12 mM sodium lauroyl sarcosinate, 100 mM triethylammonium bicarbonate buffer, 10 mM tris(2-carboxyethyl)phosphine, and 40 mM chloroacetamide was directly added to the beads, and the samples were incubated for 10 min at 95°C with shaking at 1,200 rpm. This step solubilized the EVs and denatured, reduced, and alkylated the EV proteins. After the samples were cooled down to room temperature, acetonitrile was added to the sample to a final concentration of 70% (v/v). The samples were incubated for 5 min at room temperature with shaking. The supernatants were removed, and the beads were further washed with 70% acetonitrile three times. The bead samples were resuspended in 50 μL of 100 mM triethylammonium bicarbonate buffer, and MS-grade trypsin was added at 1:50 (wt/wt) enzyme-to-protein ratio for overnight digestion at 37 °C with shaking at 1200 rpm. Acetonitrile was added to the samples to a final concentration of 60% (v/v). Peptide-containing supernatants were collected, and peptide concentration was determined using the PierceTM Quantitative Colorimetric Peptide Assay (Thermo Scientific) according to the manufacturer’s instructions. Equal amounts of peptides from each sample were aliquoted and dried in a SpeedVac.

### Stable isotope dimethyl labeling

The peptide samples were labeled with dimethyl groups using the in-solution labeling protocol (27). In brief, 25 μg of peptides were reconstituted in 100 μl of 100 mM triethylammonium bicarbonate buffer. To each dimethyl labeling reaction, 4 μl of 4% (vol/vol) formaldehyde (CH_2_O, CD_2_O, or ^13^CD_2_O) and 4 μl of 600 mM sodium cyanoborohydride (NaBH_3_CN or NaBD_3_CN) were added sequentially. The samples were incubated for 1 hr at room temperature on a bench-top incubator with shaking at 1,200 rpm. To quench the reaction, 2 μl of 10% (vol/vol) ammonia (NH_4_OH) was added to each tube on ice. The solutions were then acidified by adding 5 μl of 10% (vol/vol) formic acid (FA). The labeled peptides from different channels were pooled and desalted using in-house prepared C18 StageTips. The resulting peptides can be injected into LC-MS/MS or subjected to fractionation.

### StageTip-based high pH reverse phase fractionation

To build a project-specific spectral library, 10 μg of pooled labeled peptides were fractionated using a high-pH reverse phase condition. Dried peptides were resuspended in 100 μL of 200 mM ammonium formate (pH 10.0). In-house prepared C18 StageTips were first activated by acetonitrile and equilibrated by ammonium formate. The peptides were loaded into the tips and eluted into eight fractions with the step acetonitrile (4-80%) in a high-pH condition. The fractions were dried in a SpeedVac for LC-MS/MS analysis.

### LC-MS/MS analysis

Dried peptides were dissolved in 0.1% FA and loaded onto Evotips Pure (EvoSep) according to the manufacturer’s instructions. Peptides were then analyzed using the Evosep One LC system (EvoSep) coupled with a timsTOF HT mass spectrometer (Bruker). The 60 SPD method was used with the PepSep C18 column (8 cm x 150 μm, 1.5 μm; Bruker). The columns were connected to a ZDV Sprayer 20 μm emitter (Bruker) inside a Captive Spray source (Bruker). The mobile phases included 0.1% FA in LC–MS grade water as solvent A and 0.1% FA in LC–MS grade acetonitrile as solvent B. The TIMS dimension was calibrated using three Agilent ESI tuning mix ions (622.0289, 0.9848 V/cm^2^; 922.0097, 1.1895 V/cm^2^; 1221.9906, 1.3820 V/cm^2^) in positive mode.

For generating the spectral library and DDA analysis, DDA-PASEF mode was used to perform data acquisition of the 8 fraction samples on the timsTOF HT. Ten PASEF/MSMS scans were acquired per cycle using the 60 SPD method. Spectra were acquired within an m/z range from 100 to 1,700 and ion mobility (IM) from 0.6 to 1.6 V/cm^2^. The ion mobility-dependent collision energy was set from 59 eV at 1.6 V/cm^2^ to 20 eV at 0.6 V/cm^2^. Singly charged precursors were excluded by their position in the m/z-IM plane, and a precursor signal intensity threshold of 2,500 was used for fragmentation. Precursors were isolated with a 2 Th window (< 700 Da) and a 3 Th window (> 800 Da) and were excluded for 0.4 min when they were over a target intensity of 20,000.

The dia-PASEF methods for mDIA analysis were optimized with py_diAID using the DDA-based library and consisted of variable isolation window widths based on the precursor ion densities (43). The isolation windows covered an m/z range from 300–1450, and the IM was set from 0.7 to 1.3 V/cm^2^. The dia-PASEF method contained 12 dia-PASEF scans with a cycle time of 1.38 s. The ramp time was set as 100 ms, and the collision energy was set from 59 eV at 1.6 V/cm^2^ to 20 eV at 0.6 V/cm^2^ (see **Supplemental Table S1-S3** and **Figure S1** for detailed window placements).

### Raw data processing

The project-specific spectral library for mDIA analysis was generated by Spectronaut^TM^ v19 software (Biognosys). The input raw files included 8 DDA runs and were searched against the human database. In Pulsar search engine, trypsin/P as specific digest type, 7 minimal peptide length, 52 maximum peptide length, carbamidomethyl at cysteine as fixed modification, acetylation at protein N-term, oxidation at methionine, and 5 as maximum variable modifications were applied. Three labeling channels were enabled, including channel 1(Light, DimethLys0 and DimethNter0), channel 2 (Medium, DimethLys4 and DimethNter4), and channel 3 (Heavy, DimethLys8 and DimethNter8). The FDR at PSM, peptide, and protein group were set to 0.01. In-silico generate missing channels option was enabled.

For mDIA using Spectronaut, the dia-PASEF raw file of each sample was searched using the project-specific library or directDIA (library-free). In the library-based pipeline, retention time prediction with local (non-linear) regression calibration was used. The identification was determined with mutated decoy generation and dynamic size at 0.1 fractions of library size. The single hit definition was set to stripped sequence. The quantification was performed at the MS2 level, and cross-run normalization was enabled. Qvalue and no imputing were used as data filtering. In the library-free pipeline, directDIA+(Deep) was selected. Other settings of the Pulsar search engine and DIA analysis were the same as the above-mentioned library-based DIA settings.

For mDIA using DIA-NN (version 1.9) (44), the dia-PASEF raw file of each sample was searched using the project-specific library generated from Spectronaut or the predicted library generated from the previous study using AlphaPeptDeep Python framework (33). Since DIA-NN generates the library for the other channels on its own by the commands, we created a separate Spectronaut library containing only the Light channel. After exporting the library as a tsv file, we used the Pandas data frame library in Python 3.8.8 to rewrite the file, replacing all dimethyl labels with “[Dimethyl]” to maintain consistency with the format required for DIA-NN input. The DIA-NN search included the following settings: two missed cleavages, a maximum of 5 variable modifications per peptide, carbamidomethylation of cysteines as fixed modification, protein N-terminus acetylation and oxidation of methionine as variable modifications, protein inference = ‘Genes,’ neural network classifier = ‘Single-pass mode,’ quantification strategy = ‘Robust LC (high precision),’ cross-run normalization = ‘RT-dependent,’ library Generation = ‘IDs, RT and IM Profiling,’ and speed and RAM usage = ‘Optimal results.’ Mass accuracy and MS1 accuracy were set to 0 for automatic inference from the first run. The following settings were also enabled: ‘Use isotopologues,’ ‘MBR,’ ‘Heuristic protein inference,’ and ‘No shared spectra.’ For mDIA samples, the following additional commands were entered into the DIA-NN command line GUI: (1) {--fixed-mod Dimethyl, 28.0313, nK}, (2) {--channels Dimethyl, 0, nK, 0:0; Dimethyl, 4, nK, 4.0251:4.0251; Dimethyl, 8, nK, 8.0444:8.0444}, (3) {--original-mods}, (4) {--peak-translation}, (5) {--ms1-isotope-quant}, (6) {--report-lib-info}, and (7) {-mass-acc-quant 10.0}. DIA-NN outputs were further processed with the DIA-NN R package to perform FDR filtering (FDR ≤ 0.01). Peptides and protein groups were quantified using the MaxLFQ algorithm.

For DDA using FagPipe (version 22.0) (45), the DDA-PASEF raw file of each sample was searched using the settings in the following configuration: MSFragger (v4.1) was used to conduct a closed search followed by labeling quantification via IonQuant (v1.10.27). Both precursor and fragment mass tolerances were set to 20 ppm for the closed search. The enzyme specificity was set to strict trypsin, allowing a maximum of two missed cleavages. Methionine oxidation and protein N-terminal acetylation were included as variable modifications, while cysteine carbamidomethylation was set as a fixed modification. Dimethylation labeling (light: +28.0313, medium: +32.0564, heavy: +36.0757) on peptide N-termini and lysine residues was specified as a variable modification. Following the closed search, MSBooster, Percolator, and Philosopher were used to generate deep-learning scores, perform rescoring, and estimate the FDR for PSMs. For labeling quantification with IonQuant, the mass tolerance was set to 10 ppm, and the retention time tolerance was 0.4 min. Match-between-runs and intensity normalization across runs were enabled.

### Bioinformatics analysis

The ratio accuracy and coefficient of variation were calculated and plotted in Python (version 3.8.8) using Numpy, Pandas Dataframes, and Plotly libraries. To perform differential analysis, the data were analyzed in the Perseus software (version 2.0.7.0) using the intensities of proteins extracted from the Spectronaut or DIA-NN search results. The data were first log2-transformed, and quantifiable proteins were selected as identified three replicates in at least one condition. The imputation was performed by replacing the missing values of intensities with a normal distribution with a downshift of 1.8 SDs and a width of 0.3 SDs. The imputed data were further normalized by subtracting the median of all intensities in each individual sample from all the intensities. In the scatter plots, the significantly changed proteins were identified by the p-value obtained from a two-sample t-test (< 0.05) and fold-change (> 2 or < -2). Hierarchical clustering analysis was performed using Euclidean distance and average linkage. Visualizations of scatter plots, heatmaps, correlation plots, and box plots were created using R (version 4.2.2). KEGG pathway analysis and GO biological processes analysis were performed using ShinyGO (version 0.82) (46).

### Western blotting

The bead samples after EV isolation were directly boiled in LDS sample buffer and loaded on a gel. Aliquots of each sample were loaded at equivalent volumes and were separated on an SDS-PAGE gel (NuPAGE 4-12% Bis-Tris Gel, Thermo Fisher Scientific). The proteins were transferred onto a low-fluorescence PVDF membrane, and the membrane was blocked with 1% BSA in TBST for 1 hr. The membranes were then incubated with rabbit anti-CD9, rabbit anti-TSG101, rabbit anti-HSPA8, mouse anti-Alix, and mouse anti-Calnexin at a 1:5,000 ratio overnight in 1% BSA in TBST. The secondary antibodies visualizing the binding of primary antibodies were goat anti-rabbit HRP-linked IgG antibody and horse anti-mouse HRP-linked IgG antibody, incubated for 1 hr in 1% BSA in TBST. The substrates (Pierce™ ECL Western Blotting Substrate) were mixed in a 1:1 ratio and added onto the membrane. The membrane was scanned by ChemiDoc Touch Imaging System (Bio-Rad) for signal detection.

### Nanoparticle tracking analysis (NTA)

The eluted EVs were diluted to a final volume of 0.8 mL using PBS. Their concentration and size were analyzed using an NTA instrument (NanoSight LM10). Video data were collected three times, each with a duration of 30 s. The camera level and detection threshold were set to 5 and 13, respectively. NanoSight NTA 3.2 software was used for data analysis.

### Transmission electron microscopy (TEM)

The eluted EVs were dried using SpeedVac and resuspended in 100 μL of PBS. The sample was then sonicated in an ultrasonic bath for 30 sec. 10 μL of the EV sample was pipetted onto a 200-mesh copper grid with carbon-coated formvar film and incubated for 10 min. The grid was washed with distilled water and placed on 10 μL of 1% uranyl acetate for 2 min. Excess liquid was removed by blotting. Images were acquired using a single TEM instrument (Tecnai-T20) and DigitalMicrograph software.

### Experimental Design and Statistical Rationale

This study aims to evaluate different mDIA-based pipelines to identify an optimal approach for EV proteomics. Pipeline comparison experiments were conducted using EVs derived from the CCLP cell line, with all DDA and DIA pipelines performed in three technical replicates. To ensure consistency, EVs from the same batch and passage of cultured cells were used for both library generation and single runs. EV characterization experiments, including Western blotting and NTA, were also performed in three technical replicates. To assess proteome changes in ICC cells and their EVs, mDIA-based proteomic analyses were performed using three biological replicates for each condition (CCLP, RBE, or RBE+AG120). Among the conditions, IDH1 mutant cells without inhibitor treatment were treated with the same volume of DMSO as vehicle control. Statistical analysis parameters are detailed in the respective paragraphs or figure legends.

## Results

### Development of mDIA pipelines for EV proteomics

We designed an EV proteomics workflow to evaluate different mDIA pipelines that consider factors of spectral libraries and software suites (**Figure 1A**). Culture media from intrahepatic cholangiocarcinoma CCLP cells were collected, and EVs were isolated using the EVtrap beads functioned with chemical affinity groups (38, 40). To confirm successful EV isolation, we characterized the samples by detecting EV markers, size and shape using Western blot, NTA and TEM, respectively (**Figure 1B-D**). Signals for four positive markers, CD9, TSG101, HSPA8, and Alix, were enriched in the EV samples, while the negative marker, Calnexin, showed no signal (**Figure 1B**). The isolated sample exhibited a size distribution of 50–400 nm, consistent with the expected size range for EVs (**Figure 1C**). TEM images revealed that EVs isolated from the media displayed the typical cup-shaped morphology (**Figure 1D**). Based on these characterizations, we are confident that cell culture media-derived EVs were successfully isolated with minimal contamination from cells.

**Figure 1.**
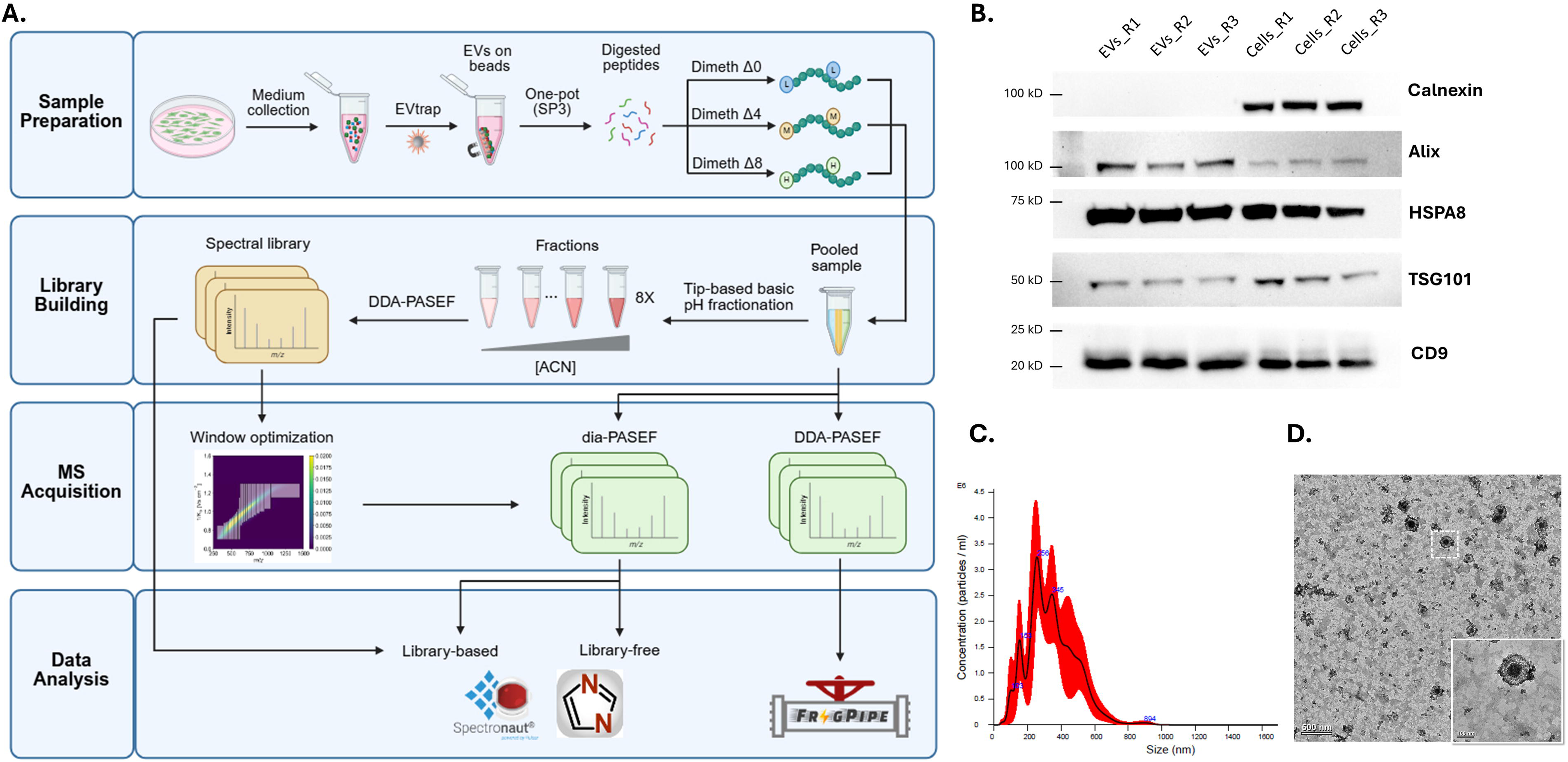
Schematic workflow of mDIA-based EV proteomics and EV characterization. A, Overview of the experimental workflow for EV proteomics using dimethyl labeling-based mDIA. EVs are isolated from conditioned media using EVtrap, followed by peptide preparation and dimethyl labeling with three isotopic channels: Dimeth Δ0 (light), Dimeth Δ4 (medium), and Dimeth Δ8 (heavy). Both DDA and DIA are used, with DIA data analyzed using library-based and library-free methods. B, Western blot analysis of EV markers. EVs and whole-cell lysates were probed for negative EV marker, Calnexin, and positive EV markers, Alix, HSPA8, TSG101, and CD9. 15 μg of lysate proteins from cells and EVs were loaded for SDS-PAGE. Each sample type was detected in triplicate. C, NTA of EVs showing the size distribution and concentration of particles in the sample. The red shading represents the standard deviation across measurements. D, TEM image of isolated EVs. The inset highlights a representative EV with characteristic morphology. Scale bar: 200 nm.

Following one-pot EV sample preparation (42), the resulting peptides were labeled separately with different dimethyl channels. For identification evaluation and building the peptide library, the three channels were pooled equally, while for quantification evaluation, they were pooled in a 1:2:4 ratio. The pooled sample was first analyzed using the conventional DDA mode, and labeling efficiency was determined. By comparing the labeled peptides relative to all detected peptides, ∼97% labeling efficiency in three channels was observed (**Supplemental Figure S2**). For library-based mDIA analysis, the pooled sample was processed with StageTip-based high-pH reverse phase fractionation, generating eight peptide fractions for the spectral library building. The resulting library contained 31,467 precursors and 6,119 proteins (**Supplemental Table S4**). This library was also analyzed using the py_diAID Python package to optimize variable isolation window widths based on precursor ion densities (**Supplemental Table S1** and **Figure S1A**) (43). The resulting optimal m/z and ion mobility window placement covered ∼99% of all precursors (**Supplemental Table S5**). This optimal dia-PASEF method was then implemented on the timsTOF platform to acquire data from single runs of the pooled sample. In the data analysis, DDA data were analyzed using FragPipe, while DIA data were analyzed using two commonly used DIA search software suites, Spectronaut and DIA-NN. In terms of library input for mDIA search, the project-specific library from the fractions was used in both Spectronuat and DIA-NN. Additionally, a predicted library from Thielert et al. was also used in DIA-NN (33). Library-free pipeline (directDIA) in Spectronaut was also included for evaluation.

### Project-specific library-based mDIA outperformed other pipelines in the identification of EV proteins

In the evaluation of protein identification (**Figure 2A and Supplemental Data S1**), DDA of EV samples without fractionation yielded ∼1,500 unique proteins based on FragPipe database search, which is comparable to previous label-free DDA results obtained from EVs in cell culture media (47, 48). With the library generated from the fractions, the protein identification number increased to 2,300-2,500 using both Spectronaut and DIA-NN. Compared to DDA, this increased identification number indicates library-based mDIA has superior sensitivity to detect the low abundant EV protein signals. When we employed directDIA in Spectronaut and a predicted library in DIA-NN to search the mDIA data, we observed a decrease in the number of identified proteins (1,400-2,100 unique proteins). This reduction may be attributed to the absence of a representative spectral library. Without accurate spectral information, such as retention time, m/z, and ion mobility for reference, fewer ions were identified, resulting in a lower overall identification number. The reproducibility of protein identification was further assessed across replicate runs for each pipeline (**Figure 2B**). The DDA pipeline demonstrated the lowest reproducibility, with an identification overlap of 68.2% across triplicates. The directDIA method also exhibited relatively poor reproducibility (66.8%), confirming the limitations of searching mDIA data without a representative spectral library. Similarly, DIA-NN using a predicted library showed a reduced overlap of 73.7%. In contrast, the mDIA pipelines using a project-specific experimental library showed higher identification reproducibility. These results demonstrate the advantage of employing experimentally derived spectral libraries for improving both identification sensitivity and reproducibility for EV samples.

**Figure 2.**
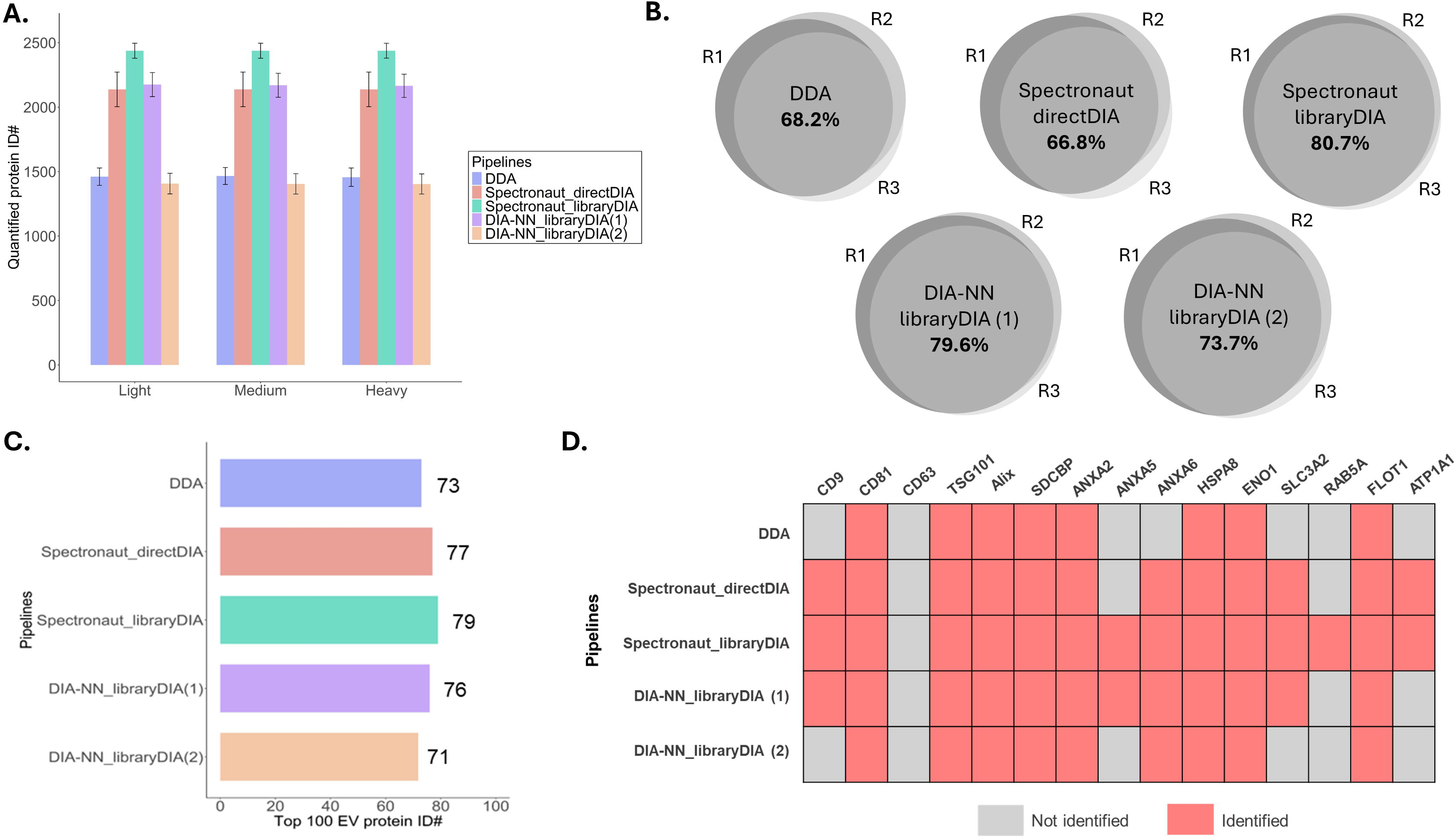
Evaluation of identification outcomes from different mDIA pipelines. A, bar chart comparing the number of quantified protein identifications across light, medium, and heavy labeling channels using different analysis pipelines. Error bars represent standard deviation across triplicates. B, Venn diagrams showing the percentage overlap of protein identifications among triplicates for each analysis pipeline. C, the number of proteins from the top 100 most common EV proteins in the ExoCarta database identified by each analysis pipeline. D, heatmap displaying the identification status of selected EV marker proteins across different analysis pipelines. Red indicates identified proteins, while gray represents proteins that were not identified. (1) project-specific fraction-based library. (2) predicted library from Thielert et al.

To further evaluate the specificity of these pipelines, we compared the identified proteins with the top 100 commonly reported EV proteins in the ExoCarta database (**Figure 2C**) (49). The DDA pipeline exhibited the lowest overlap with this list. Among all pipelines, library-DIA using Spectronaut achieved the highest overlap, further supporting the superior specificity of the library-based mDIA in detecting known EV proteins. When analyzing specific EV marker proteins (**Figure 2D**), distinct differences emerged among the pipelines. For example, CD9, a widely recognized EV marker (39), was only identified in DIA-based pipelines and was absent in DDA results. Additionally, Rab5A, a protein involved in EV trafficking (50), was only detected in the result of library-DIA using Spectronaut. These findings indicate the enhanced specificity and sensitivity of library-based mDIA pipelines for EV protein identification. Overall, our results highlight the superior proteome coverages achieved by library-based mDIA pipelines over conventional DDA and directDIA pipelines.

### Comparison of quantitative outcomes from different mDIA pipelines

Next, quantification outcomes from different mDIA pipelines were evaluated. To assess quantitative variation between channels, we input equal amounts of labeled peptides into each channel and calculated pairwise correlation coefficients across channels within each pipeline. (**Figure 3A and Supplemental Data S1**). All pipelines demonstrated high correlation values (>0.9) across channels, indicating high quantification consistency. In addition to correlation, we also evaluated the intensity distribution of identified proteins within each pipeline. The DDA pipeline showed a slightly skewed distribution, deviating from normality. This skew may be attributed to co-isolation interference, which leads to chimeric spectra and affects MS1-level quantification. In contrast, DIA-based pipelines, especially those using Spectronaut, exhibited more normally distributed intensity values, reflecting improved data quality.

**Figure 3.**
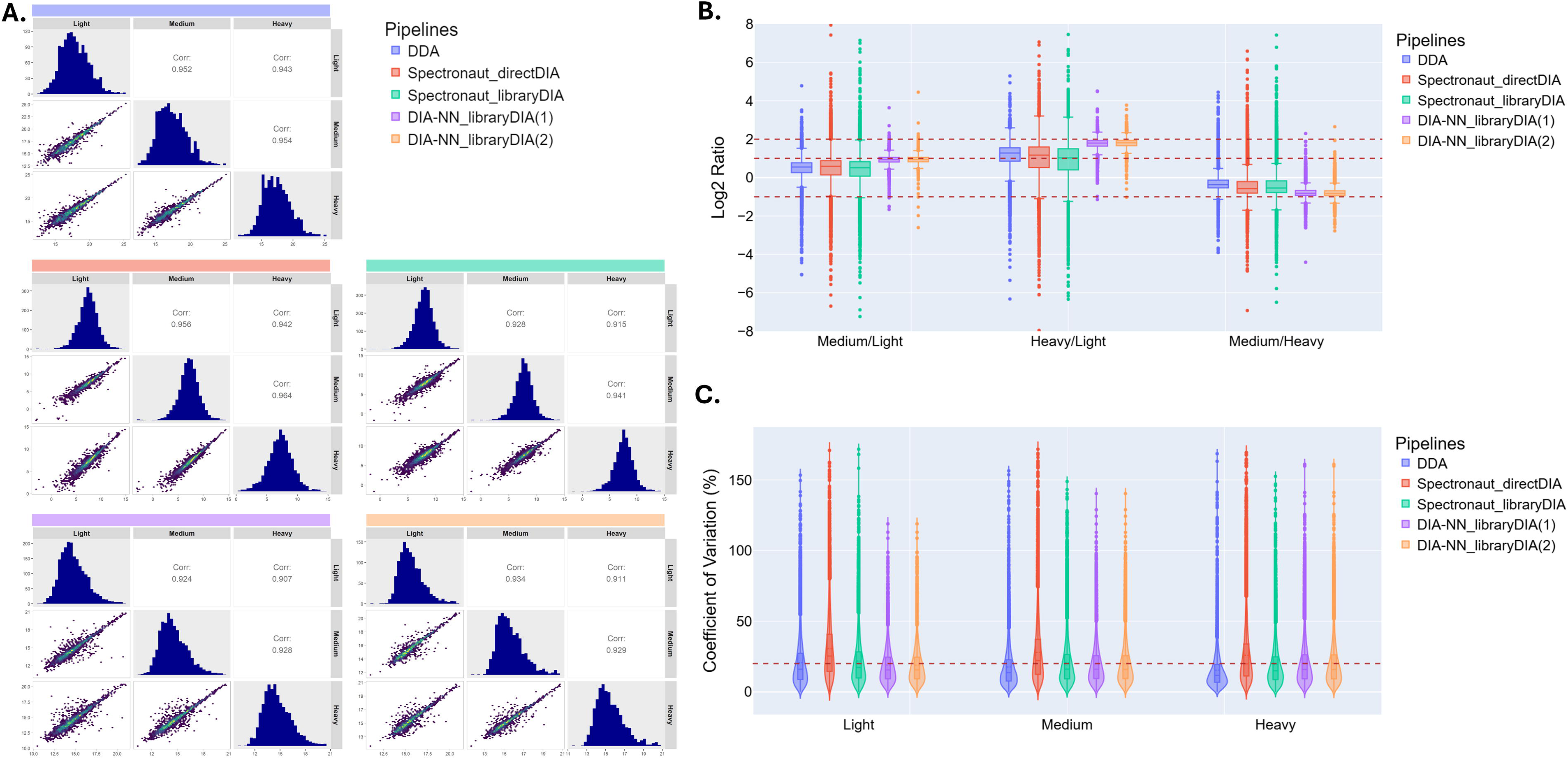
Evaluation of quantification outcomes from different mDIA pipelines. A, pairwise correlation analysis of quantified protein intensities across different labeling channels (light, medium, and heavy) for each analysis pipeline. Each panel shows histograms of the protein intensity distribution in each channel and scatter plots with Pearson correlation coefficients (Corr) between channels. B, boxplots of log₂-transformed precursor intensity ratios for medium/light, heavy/light, and medium/heavy comparisons across pipelines. The dashed red lines indicate the expected log₂ ratios. C, CV distribution for precursors quantified across different pipelines and labeling channels. Violin plots display the spread of CV%, with a red dashed line indicating a 20% CV threshold. (1) project-specific fraction-based library. (2) predicted library from Thielert et al.

To evaluate quantification accuracy, we analyzed the distribution of intensity ratios between channels (**Figure 3B and Supplemental Data S2**). A known input ratio of 1:2:4 (light/medium/heavy) was used to benchmark the accuracy of each pipeline. All pipelines achieved comparable accuracy, with distributions centered around the expected ratios. Notably, the library-based mDIA using DIA-NN exhibited slightly better accuracy, showing minimal deviation from the theoretical ratios. We further examined the reproducibility of quantification across replicates by analyzing the coefficient of variation (CV) (**Figure 3C and Supplemental Data S2**). Most pipelines achieved acceptable CV values (<20%). However, the directDIA pipeline exhibited CV values exceeding 20% in all channels, indicating less reproducible quantification than other methods. The higher variation in the directDIA pipeline suggests that the absence of a project-specific spectral library may contribute to increased quantification variability.

### ICC EV proteome reflects cellular responses to IDH1 mutation

Given the superior identification and quantification performance of the optimal mDIA pipeline using library-based DIA with Spectronaut, we further applied it to investigate EV proteome changes associated with IDH1 mutation in ICC. IDH1 plays an important role in cellular metabolism by converting isocitrate to alpha-ketoglutarate. Mutations in IDH1, frequently found in gliomas, acute myeloid leukemia, and cholangiocarcinoma, result in a gain-of-function activity that produces the oncometabolite 2-hydroxyglutarate (2-HG) (51). The accumulation of 2-HG disrupts cellular differentiation and drives tumor progression by altering gene expression and epigenetic regulation (52). Due to its specific and abnormal metabolic effects, IDH1 mutation has become an important drug target in cancer research, leading to the development of inhibitors such as AG-120 (Ivosidenib), which was recently approved for glioma treatment (53, 54). While cellular gene expression and proteomic alterations associated with IDH1 mutation in ICC have been extensively studied (55), there is a lack of studies exploring proteome changes in EVs, especially in response to inhibitor treatment. Investigating dysregulated EV proteins could provide important information on potential biomarkers for cancers with IDH1/2 mutations such as glioma where tissue extraction for diagnosis is extremely difficult and invasive.

We collected EVs from CCLP (wild-type IDH1) and RBE (mutant IDH1) cells, along with EVs from RBE cells treated with AG-120 (**Figure 4A**). EV proteins were extracted and digested into peptides. After dimethyl labeling of peptides from three conditions, we performed StageTip-based fractionation on the pooled sample and built the project-specific library for window optimization (**Supplemental Table S2 and S5** and **Figure S1B)** and mDIA using Spectronaut (**Supplemental Table S4**). This pipeline enabled the quantification of over 2,000 unique proteins (**Supplemental Data S3**) and again obtained improved quantitative precision compared to the result using directDIA (**Supplemental Figure S3**). To determine whether EVs reflect key cellular proteomic changes, we also profiled cell proteomes using DIA-NN with a predicted library, as previously described by Thielert et al (**Supplemental Data S4**) (33). A comparison of identified proteins in cell and EV samples revealed that more than 90% of EV proteins overlapped with the cellular proteome (**Figure 4B**).

**Figure 4.**
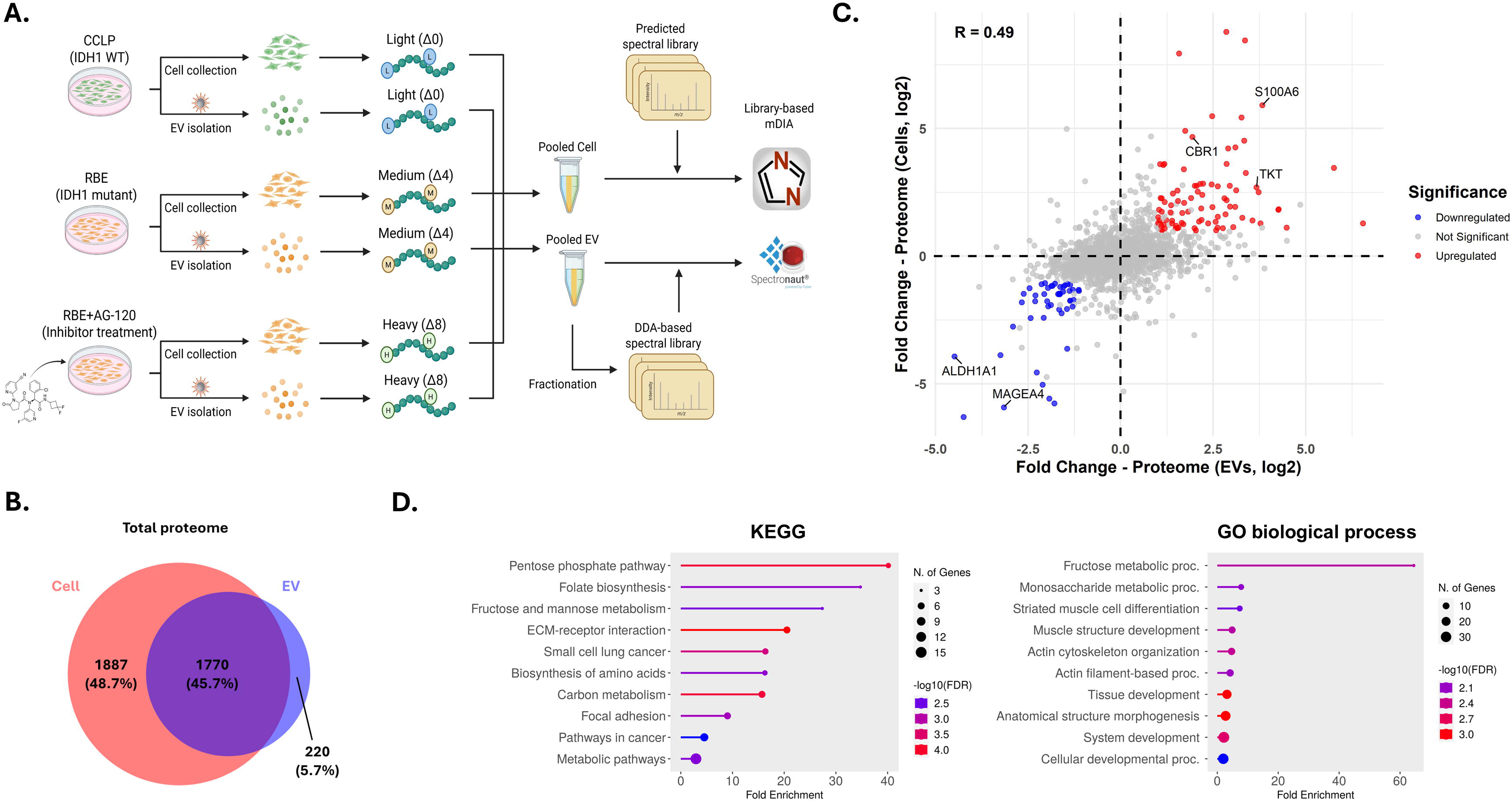
Application of the optimal DIA pipeline for detecting EV proteome changes caused by IDH1 mutation in ICC. A, Schematic of the experimental workflow for detecting both cellular and EV proteomes. Cell lines used include CCLP (IDH1 WT), RBE (IDH1 mutant), and RBE treated with AG-120 (inhibitor treatment). Cells and EVs were collected from different plates and processed using the optimal mDIA pipeline. B, Venn diagram of the total proteomes from cells and EVs, including proteins that were unique to cells (red), EVs (blue), or shared between them (purple). C, correlation of proteome changes caused by IDH1 mutation between cells and EVs. A scatter plot shows the log_2_ fold change in protein abundance in cells versus EVs. Significantly (p-value < 0.05, |fold change| > 2) upregulated proteins are shown in red, and downregulated proteins in blue. Representative proteins are labeled. The Pearson correlation coefficient is shown. D, KEGG pathway and GO biological process enrichment of upregulated proteins shared in both cells and EVs. Circle size indicates the number of genes, and color represents the significance level (-log_10_(FDR)).

We first compared the RBE condition with the CCLP condition. Interestingly, the majority of proteins in both cells and EVs exhibited similar fold changes between the two conditions (**Figure 4C**). In both EVs and cells, more than 200 proteins were significantly up-or down-regulated in RBE conditions (**Supplemental Figure S5A and S5C**). Among them, we identified 84 upregulated and 48 downregulated proteins shared between both sample types (**Figure 4C**), including multiple proteins, such as CBR1, S100A6, TKT, ALDH1A1, and MAGEA4, previously implicated in IDH1 mutation (56–60). To gain further insight into the functional implications of these proteomic alterations, we performed KEGG and gene ontology (GO) enrichment analyses on the shared upregulated proteins (**Figure 4D**). Several known pathways associated with IDH1 mutation, such as the pentose phosphate and monosaccharide metabolism pathways, were significantly enriched (61, 62). These findings suggest that EV proteomes indeed reflect key cellular pathways altered by IDH1 mutations.

### EV proteins as indicators of IDH1 mutant inhibitor treatment

To investigate the impact of mutant IDH1 inhibition on the EV proteome, we further compared EVs from RBE cells with and without AG-120 treatment. A total of 63 proteins were found to be significantly regulated upon AG-120 treatment (**Figure 5A**). Among these proteins, SNRPB and GRHPR were previously reported to be associated with IDH1 mutations, suggesting potential as indicators of treatment response in IDH1-mutant ICC (63, 64). Additionally, multiple proteins associated with RNA metabolism and translation were significantly downregulated in the AG-120-treated group. GO enrichment analysis further revealed significant downregulation of pathways associated with RNA capping, hypermethylation, and spliceosome function (**Figure 5B**). RNA capping and splicing are essential for mRNA maturation and translation, and their disruption by IDH mutations alters global gene expression (65). The downregulation of these pathways following AG-120 treatment might suggest a shift toward normal RNA processing in the IDH1 mutant cells. We next compared the significantly regulated proteins between EV and cell proteomes after AG-120 treatment (**Supplemental Figure S5B and S5D**) and found that only a few proteins such as PRPF4 and GLG1 were commonly regulated between cells and EVs (**Figure 5C**). The spliceosome component PRPF4, which plays an important role in mRNA splicing, was significantly upregulated in both cells and EVs (66). This again suggests that the inhibitor may affect pathways related to mRNA processing. However, the low degree of overlap suggests that the mechanisms driving proteome alterations in EVs following inhibitor treatment may not simply mirror those occurring in cells. The molecular cargo affected by the inhibitor could be selectively packaged into EVs, rather than directly reflecting proportional changes within the parent cells (67, 68). Interestingly, several proteins involved in oxidative stress responses, such as GSS (glutathione synthetase), were significantly regulated in cells, consistent with previous reports that AG-120 disrupts glutamine metabolism and increases oxidative stress (69). This cellular response, however, does not appear to be reflected in the EV proteome.

**Figure 5.**
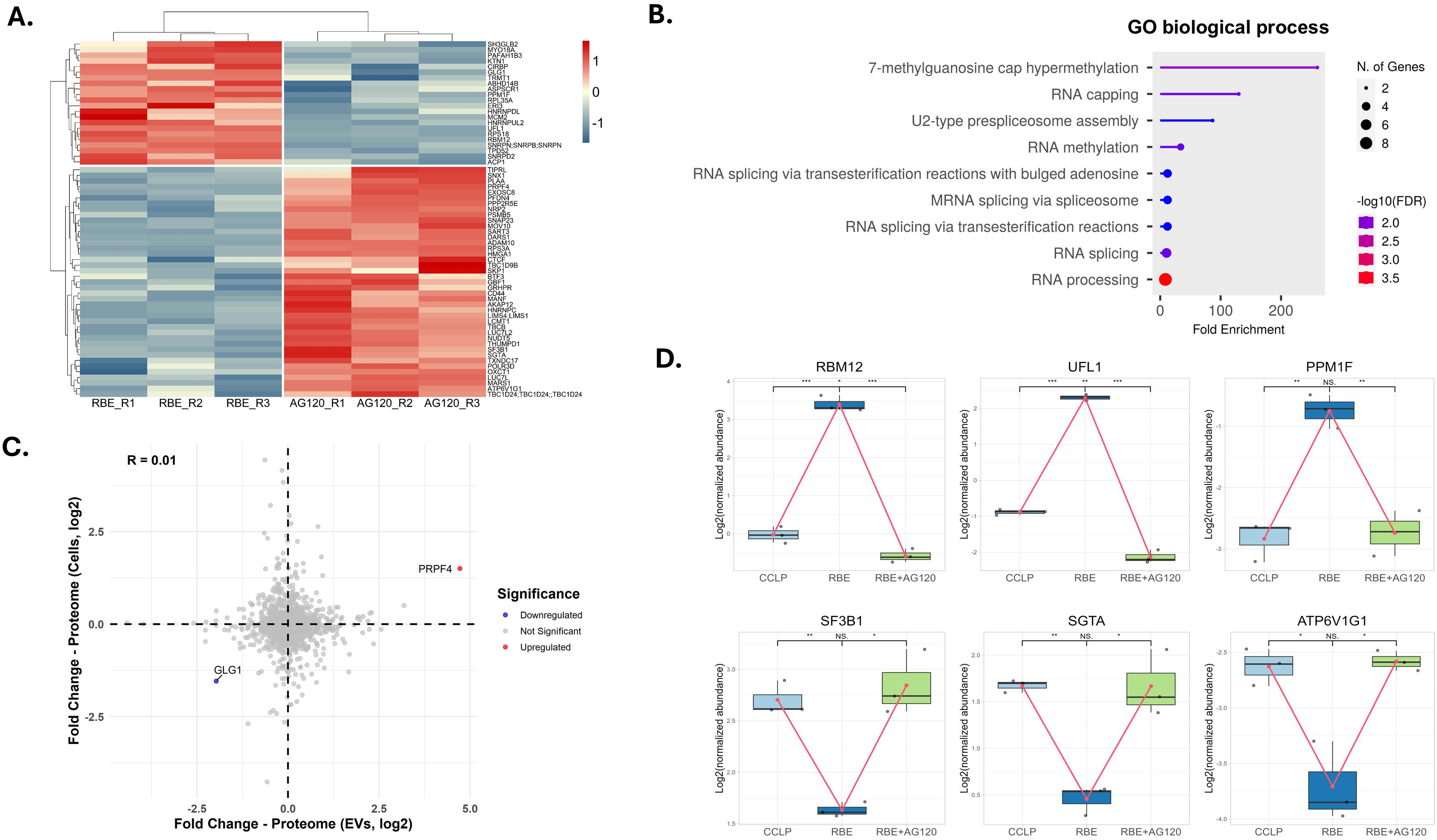
EV proteome changes caused by inhibitor treatment. A, heatmap of significantly regulated (p-value < 0.05, |fold change| > 2) proteins in EVs after AG-120 treatment. Normalized protein levels are shown for RBE (IDH1 mutant) and RBE treated with AG-120 (inhibitor treatment). Hierarchical clustering was performed, and the color scale represents Z-scores, with red indicating higher expression and blue indicating lower expression. B, GO biological process enrichment analysis of significantly downregulated EV proteins. C, Correlation of proteome changes caused by AG-120 treatment between cells and EVs. A scatter plot shows the log_2_ fold change in protein abundance in cells versus EVs. Significantly (p-value < 0.05, |fold change| > 2) upregulated proteins are shown in red, and downregulated proteins in blue. All significant proteins are labeled. The Pearson correlation coefficient is shown. D, box plots of selected differentially regulated EV proteins. Expression levels of RBM12, UFL1, PPM1F, SF3B1, SGTA, and ATP6V1G1 across CCLP (IDH1 WT), RBE (IDH1 mutant), and RBE treated with AG-120 are shown. Box plots display median, interquartile range, and individual data points. Statistical significance is indicated (*p-value < 0.05, ** p-value < 0.01, *** p-value < 0.001).

To identify potential EV proteins that reflect IDH1 mutation status in ICC and its response to inhibition, we specifically examined proteins that exhibited opposite expression patterns between the IDH1 mutant and AG-120 treatment conditions (**Supplemental Figure S6**). These proteins were significantly upregulated in EVs from IDH1 mutant cells but significantly downregulated following AG-120 treatment, or vice versa. The rationale for focusing on these proteins is based on the hypothesis that effective inhibition of mutant IDH1 should reverse the proteomic signature in EVs. For instance, proteins such as RBM12, UFL1, PPM1F, SF3B1, SGTA, and ATP6V1G1 followed this pattern (**Figure 5D**). Consistent with our findings in EVs, UFL1 has been found to exhibit higher expression levels in formalin-fixed, paraffin-embedded (FFPE) glioma samples from IDH-mutant patients compared to IDH-wild-type samples (70). Additionally, ATP6V1G1 has been reported to be downregulated in primary cells derived from IDH1/2-mutant glioma patients relative to those from IDH1/2 wild-type individuals (71). Taken altogether, the presence of these proteins in EVs may serve as a potential indicator for identifying IDH1 mutant ICC subtype and monitoring AG-120 therapeutic response.

## Discussion

The success of EV proteome profiling requires an efficient EV enrichment technique, an optimized sample preparation, and advanced MS data acquisition and quantitation. Expanding EV proteomics for large-scale studies or clinical applications involves higher throughput while maintaining comprehensive proteome coverage, high EV specificity, and reliable quantification. In this study, we established an optimized mDIA pipeline to address these challenges, providing a robust and effective tool for EV proteomics.

The DIA method has been extensively studied, with multiple pipelines benchmarked and compared. However, few studies have evaluated the impact of different pipelines on mDIA, which presents further spectral complexity. Most mDIA studies rely on either spectrum-centric or prediction-based library-free analysis (32, 33), but the relative performance of empirical spectral libraries remains unclear. Given their distinct nature compared to cellular proteomes, EV proteomes are less complex but primarily consist of low-abundance proteins. Additionally, depending on the EV source and isolation technique, they may also contain high-abundance proteins from contaminants. Therefore, we believe that the representativeness of the spectral library is the most critical factor in determining mDIA outcomes, particularly for EV-specific protein identification and quantification. We hypothesized that a low-capacity StageTip-based fractionation could generate sufficient fractions for building an in-depth and representative spectral library for multiplexed EV proteomes. Our results supported this hypothesis, demonstrating that library-based mDIA pipelines provided superior identification and quantification compared to library-free pipelines (**Figures 2 and 3**). However, this advantage did not extend to cell samples, where using a predicted library achieved better coverage (**Supplemental Figure S4**). This suggests that tip-based fractionation may not sufficiently resolve the complexity of multiplexed cell samples, necessitating more extensive offline liquid chromatography fractionation for in-depth proteome coverage and improved quantification. Collectively, our findings underscore the value of utilizing small-scale tip-based fractionation for library-based mDIA analysis of EV proteome.

Intrahepatic cholangiocarcinoma (ICC) is a rare but aggressive malignancy, with the IDH1 mutant subtype representing a clinically relevant subgroup characterized by poor prognosis, lower overall survival rates, and a high risk of recurrence (72). Patients with this subtype often exhibit resistance to conventional therapies, posing significant challenges in treatment and disease management. Identifying IDH1 mutant subtype is essential for guiding targeted therapeutic interventions, such as AG-120 treatment. EVs are promising candidates for biomarker discovery and non-invasive liquid biopsy applications. However, there is a lack of studies identifying EV protein markers for ICC diagnosis and treatment monitoring. In this study, we provide the first report of EV protein signatures reflecting IDH1 mutation-associated cellular pathway alterations and responses to AG-120 treatment (**Figures 4 and 5**). Although several known IDH1 mutation-associated proteins and pathways were significantly regulated in both RBE cells and their EVs, the overlap of significantly regulated proteins between cells and EVs in the AG-120 treatment condition was limited (**Figure 5C**). This may suggest that EVs selectively package certain proteins in response to AG-120 treatment rather than merely reflecting intracellular proteomic changes. However, the EV proteome changes in the AG-120 treatment condition are still aligned with previous studies showing that IDH1 mutations influence epigenetic regulation, including modifications in RNA methylation and processing. RNA capping and hypermethylation are crucial for mRNA stability and translation initiation, shaping global gene expression programs (73, 74). The observed downregulation of these pathways suggests that AG-120 treatment may restore a more physiologically normal RNA processing state, potentially counteracting the oncogenic effects of mutant IDH1. However, none of the downregulated proteins identified have been previously linked to AG-120 treatment, emphasizing the need for further validation to determine their functional significance in IDH1-mutant ICC. Our study also identifies candidate EV proteins that warrant further investigation as potential biomarkers for IDH1 mutant subtype characterization and AG-120 treatment response assessment (**Figure 5D**). While more studies and validation are necessary, our findings demonstrate the potential of leveraging mDIA-based EV proteomics for biomarker discovery in cancer diagnosis and treatment monitoring.

For the clinical application of EV proteomics, achieving higher throughput is essential. Our established pipeline, which incorporates EVtrap and mDIA, is well-suited for integration with automated sample preparation, thereby facilitating scalability for clinical studies (75). However, further evaluation of the mDIA approach using clinically relevant samples, such as plasma-derived EVs, is required to assess its translational potential. Additionally, for large-scale clinical sample analyses, incorporating a bridge or reference channel is recommended to minimize batch effects and enhance quantitative performance (33, 76).

In conclusion, this study establishes a robust mDIA pipeline for quantitative EV proteomics, incorporating dimethyl labeling and project-specific spectral libraries generated through StageTip-based fractionation. Using this optimized workflow, we successfully identified proteomic alterations in EVs derived from IDH1-mutant ICC cells, highlighting changes in key metabolic pathways. Furthermore, we identified EV proteins responsive to AG-120 treatment, emphasizing the potential of EV-based biomarkers for both diagnostic and therapeutic monitoring. Our findings demonstrate the value of mDIA in advancing EV biomarker discovery, laying the groundwork for future studies focused on developing non-invasive biomarker panels for disease diagnosis and treatment monitoring.

## Data Availability

All mass spectrometry data, including raw files and search results, have been deposited to the ProteomeXchange Consortium (http://proteomecentral.proteomexchange.org) via the jPOST partner repository with the dataset identifier PXD063617.

## Supplemental data

This article contains supplemental data.

## Acknowledgments

The authors would like to thank Dr. Anton Iliuk from Tymora Analytical Operations for providing the EVtrap beads used in this study. We also thank Dr. Meng-Ju Wu from UMass Chan Medical School for generously providing the cell lines used in this study. Special thanks to Dr. Jonathan Shannahan for generously providing access to the NTA instrument.

## Author contributions

Y.-K.L., W.A.T. conceptualized the study. Y.-K.L., M.H., Z.Z., W.A.T. designed experiments. Y.-K.L., N.M. performed experiments and data analyses. Y.-K.L., N.M. generated figures and tables and wrote the manuscript with input from all authors. All authors reviewed and approved the manuscript.

## Funding and additional information

This work was partly funded by the NIH grants 4R01AG064250 (to W.A.T.) and P30 CA023168 (to Purdue Institute for Cancer Research), and by National Science Foundation grant CHE-2404098. The graphical abstract and workflow illustrations were created with BioRender.com.

## Abbreviations

EV: Extracellular vesicle
MS: Mass spectrometry
DDA: Data-dependent acquisition
mDIA: Multiplexed data-independent acquisition
PASEF: Parallel Accumulation Serial Fragmentation
FDR: False discovery rate
PSM: Peptide-spectrum match
EVtrap: Extracellular vesicle total recovery and purification
StageTip: Stop and go extraction tip
ICC: Intrahepatic cholangiocarcinoma
IDH1: Isocitrate dehydrogenase 1
AG-120: Ivosidenib
NTA: Nanoparticle tracking analysis
TEM: Transmission electron microscopy
KEGG: Kyoto Encyclopedia of Genes and Genomes

