## Supplemental Tables and Figures for "Multiplexed Data-Independent Acquisition (mDIA) to Profile Extracellular Vesicle Proteomes"

### Quantitative Extracellular Vesicle Proteomics by Multiplexed Data-Independent Acquisition

#### Supplemental Data

Table S1. Variable window sizes for the optimal dia-PASEF method used in the benchmarking experiment (CCLP, EVs).

Table S2. Variable window sizes for the optimal dia-PASEF method used in the application experiment (CCLP+RBE+AG120, EVs).

Table S3. Variable window sizes for the optimal dia-PASEF method used in the application experiment (CCLP+RBE+AG120, cells).

Table S4. DDA-based spectral libraries generated from fractions.

Table S5. Evaluation of optimized dia-PASEF window placement using spectral libraries.

Figure S1. Optimal dia-PASEF acquisition schemes for dimethyl labeling based mDIA.

Figure S2. Dimethyl labeling efficiency.

Figure S3. Comparison of quantification precision between library-based and library-free pipelines in the EV-based application experiment.

Figure S4. Protein identification numbers from the cell-based application experiment using different mDIA pipelines.

Figure S5. Significantly regulated proteins in cell- and EV-based experiments.

Figure S6. EV protein candidates for identifying IDH1 mutant iCCA subtype and monitoring AG-120 therapeutic response.

**Supplemental Table S1. Variable window sizes for the optimal dia-PASEF method used in the benchmarking experiment (CCLP, EVs).**

| MS Type | Cycle Id | 1/K0 Begin [Vs/cm2] | 1/K0 End [Vs/cm2] | Start Mass [m/z] | End Mass [m/z] | CE [eV] |
|---|---|---|---|---|---|---|
| MS1 | 0 | - | - | - | - | - |
| PASEF | 1 | 0.85 | 1.3 | 624.84 | 646.71 | - |
| PASEF | 1 | 0.7 | 0.85 | 300.19 | 390.93 | - |
| PASEF | 2 | 0.88 | 1.3 | 646.71 | 669.09 | - |
| PASEF | 2 | 0.7 | 0.88 | 390.93 | 423.62 | - |
| PASEF | 3 | 0.89 | 1.3 | 669.09 | 693.89 | - |
| PASEF | 3 | 0.7 | 0.89 | 423.62 | 449.29 | - |
| PASEF | 4 | 0.9 | 1.3 | 693.89 | 720.37 | - |
| PASEF | 4 | 0.7 | 0.9 | 449.29 | 469.55 | - |
| PASEF | 5 | 0.92 | 1.3 | 720.37 | 747.4 | - |
| PASEF | 5 | 0.7 | 0.92 | 469.55 | 488.74 | - |
| PASEF | 6 | 0.93 | 1.3 | 747.4 | 777.39 | - |
| PASEF | 6 | 0.7 | 0.93 | 488.74 | 507.95 | - |
| PASEF | 7 | 0.94 | 1.3 | 777.39 | 808.46 | - |
| PASEF | 7 | 0.7 | 0.94 | 507.95 | 527.32 | - |
| PASEF | 8 | 0.96 | 1.3 | 808.46 | 846.47 | - |
| PASEF | 8 | 0.7 | 0.96 | 527.32 | 545.81 | - |
| PASEF | 9 | 0.98 | 1.3 | 846.47 | 892.91 | - |
| PASEF | 9 | 0.7 | 0.98 | 545.81 | 564.83 | - |
| PASEF | 10 | 1 | 1.3 | 892.91 | 956.06 | - |
| PASEF | 10 | 0.7 | 1 | 564.83 | 584.55 | - |
| PASEF | 11 | 1.03 | 1.3 | 956.06 | 1045.6 | - |
| PASEF | 11 | 0.7 | 1.03 | 584.55 | 603.83 | - |
| PASEF | 12 | 1.15 | 1.3 | 1045.6 | 1444.23 | - |
| PASEF | 12 | 0.7 | 1.15 | 603.83 | 624.84 | - |

**Supplemental Table S2. Variable window sizes for the optimal dia-PASEF method used in the application experiment (CCLP+RBE+AG120, EVs).**

| MS Type | Cycle Id | 1/K0 Begin [Vs/cm2] | 1/K0 End [Vs/cm2] | Start Mass [m/z] | End Mass [m/z] | CE [eV] |
|---|---|---|---|---|---|---|
| MS1 | 0 | - | - | - | - | - |
| PASEF | 1 | 0.86 | 1.3 | 647.89 | 670.39 | - |
| PASEF | 1 | 0.7 | 0.86 | 300.52 | 395.55 | - |
| PASEF | 2 | 0.89 | 1.3 | 670.39 | 694.06 | - |
| PASEF | 2 | 0.7 | 0.89 | 395.55 | 433.92 | - |
| PASEF | 3 | 0.91 | 1.3 | 694.06 | 719.36 | - |
| PASEF | 3 | 0.7 | 0.91 | 433.92 | 461.79 | - |
| PASEF | 4 | 0.92 | 1.3 | 719.36 | 746.88 | - |
| PASEF | 4 | 0.7 | 0.92 | 461.79 | 485.5 | - |
| PASEF | 5 | 0.93 | 1.3 | 746.88 | 774.39 | - |
| PASEF | 5 | 0.7 | 0.93 | 485.5 | 506.28 | - |
| PASEF | 6 | 0.95 | 1.3 | 774.39 | 805.39 | - |
| PASEF | 6 | 0.7 | 0.95 | 506.28 | 526.27 | - |
| PASEF | 7 | 0.96 | 1.3 | 805.39 | 838.95 | - |
| PASEF | 7 | 0.7 | 0.96 | 526.27 | 545.95 | - |
| PASEF | 8 | 0.98 | 1.3 | 838.95 | 877.51 | - |
| PASEF | 8 | 0.7 | 0.98 | 545.95 | 565.82 | - |
| PASEF | 9 | 1 | 1.3 | 877.51 | 924.01 | - |
| PASEF | 9 | 0.7 | 1 | 565.82 | 585.6 | - |
| PASEF | 10 | 1.02 | 1.3 | 924.01 | 985.54 | - |
| PASEF | 10 | 0.7 | 1.02 | 585.6 | 605.35 | - |
| PASEF | 11 | 1.05 | 1.3 | 985.54 | 1070.57 | - |
| PASEF | 11 | 0.7 | 1.05 | 605.35 | 626.35 | - |
| PASEF | 12 | 1.15 | 1.3 | 1070.57 | 1398.93 | - |
| PASEF | 12 | 0.7 | 1.15 | 626.35 | 647.89 | - |

**Supplemental Table S3. Variable window sizes for the optimal dia-PASEF method used in the application experiment (CCLP+RBE+AG120, cells).**

| MS Type | Cycle Id | 1/K0 Begin [Vs/cm2] | 1/K0 End [Vs/cm2] | Start Mass [m/z] | End Mass [m/z] | CE [eV] |
|---|---|---|---|---|---|---|
| MS1 | 0 | - | - | - | - | - |
| PASEF | 1 | 0.85 | 1.3 | 638.41 | 660.86 | - |
| PASEF | 1 | 0.7 | 0.85 | 300.84 | 381.55 | - |
| PASEF | 2 | 0.88 | 1.3 | 660.86 | 683.89 | - |
| PASEF | 2 | 0.7 | 0.88 | 381.55 | 418.9 | - |
| PASEF | 3 | 0.89 | 1.3 | 683.89 | 709.03 | - |
| PASEF | 3 | 0.7 | 0.89 | 418.9 | 447.73 | - |
| PASEF | 4 | 0.91 | 1.3 | 709.03 | 736.43 | - |
| PASEF | 4 | 0.7 | 0.91 | 447.73 | 471.25 | - |
| PASEF | 5 | 0.92 | 1.3 | 736.43 | 765 | - |
| PASEF | 5 | 0.7 | 0.92 | 471.25 | 492.94 | - |
| PASEF | 6 | 0.94 | 1.3 | 765 | 796.41 | - |
| PASEF | 6 | 0.7 | 0.94 | 492.94 | 513.81 | - |
| PASEF | 7 | 0.96 | 1.3 | 796.41 | 830.42 | - |
| PASEF | 7 | 0.7 | 0.96 | 513.81 | 534.29 | - |
| PASEF | 8 | 0.98 | 1.3 | 830.42 | 868.41 | - |
| PASEF | 8 | 0.7 | 0.98 | 534.29 | 555.32 | - |
| PASEF | 9 | 1 | 1.3 | 868.41 | 914.77 | - |
| PASEF | 9 | 0.7 | 1 | 555.32 | 575.82 | - |
| PASEF | 10 | 1.03 | 1.3 | 914.77 | 977.89 | - |
| PASEF | 10 | 0.7 | 1.03 | 575.82 | 595.86 | - |
| PASEF | 11 | 1.06 | 1.3 | 977.89 | 1063.98 | - |
| PASEF | 11 | 0.7 | 1.06 | 595.86 | 616.04 | - |
| PASEF | 12 | 1.2 | 1.3 | 1063.98 | 1398.93 | - |
| PASEF | 12 | 0.7 | 1.2 | 616.04 | 638.41 | - |

**Supplemental Table S4**. **DDA-based spectral libraries generated from fractions.**

| Libraries | No. of precursors | No. of modified peptides | No. of peptides | No. of proteins |
|---|---|---|---|---|
| EVs_Benchmarking (CCLP) | 31,467 | 27,126 | 26,003 | 6,119 |
| EVs_Application (CCLP+RBE+AG120) | 48,401 | 42,625 | 38,860 | 9,410 |
| Cells_Application (CCLP+RBE+AG120) | 46,054 | 38,095 | 35,732 | 7,487 |

**Supplemental Table S5**. **Evaluation of optimized dia-PASEF window placement using spectral libraries.**

| Evaluation parameters | EVs_Benchmarking (CCLP) | EVs_Application (CCLP+RBE+AG120) | Cells_Application (CCLP+RBE+AG120) |
|---|---|---|---|
| Precursors within m/z-range [%] | 99.98 | 99.96 | 99.95 |
| Smallest diaPASEF window | 18.49 | 19.68 | 20.04 |
| Biggest diaPASEF window | 398.63 | 328.36 | 334.95 |
| Average diaPASEF window size | 47.67 | 45.77 | 45.75 |
| All proteins covered | 99.90% | 99.90% | 100.00% |
| All precursors covered | 98.60% | 98.70% | 98.10% |
| All doubly charged precursors covered | 100.00% | 100.00% | 99.90% |
| All triply charged precursors covered | 96.20% | 95.40% | 94.90% |
| All quadruply charged precursors covered | 99.40% | 99.70% | 98.60% |
| All singly charged precursors covered | 100.00% | 100.00% | 100.00% |

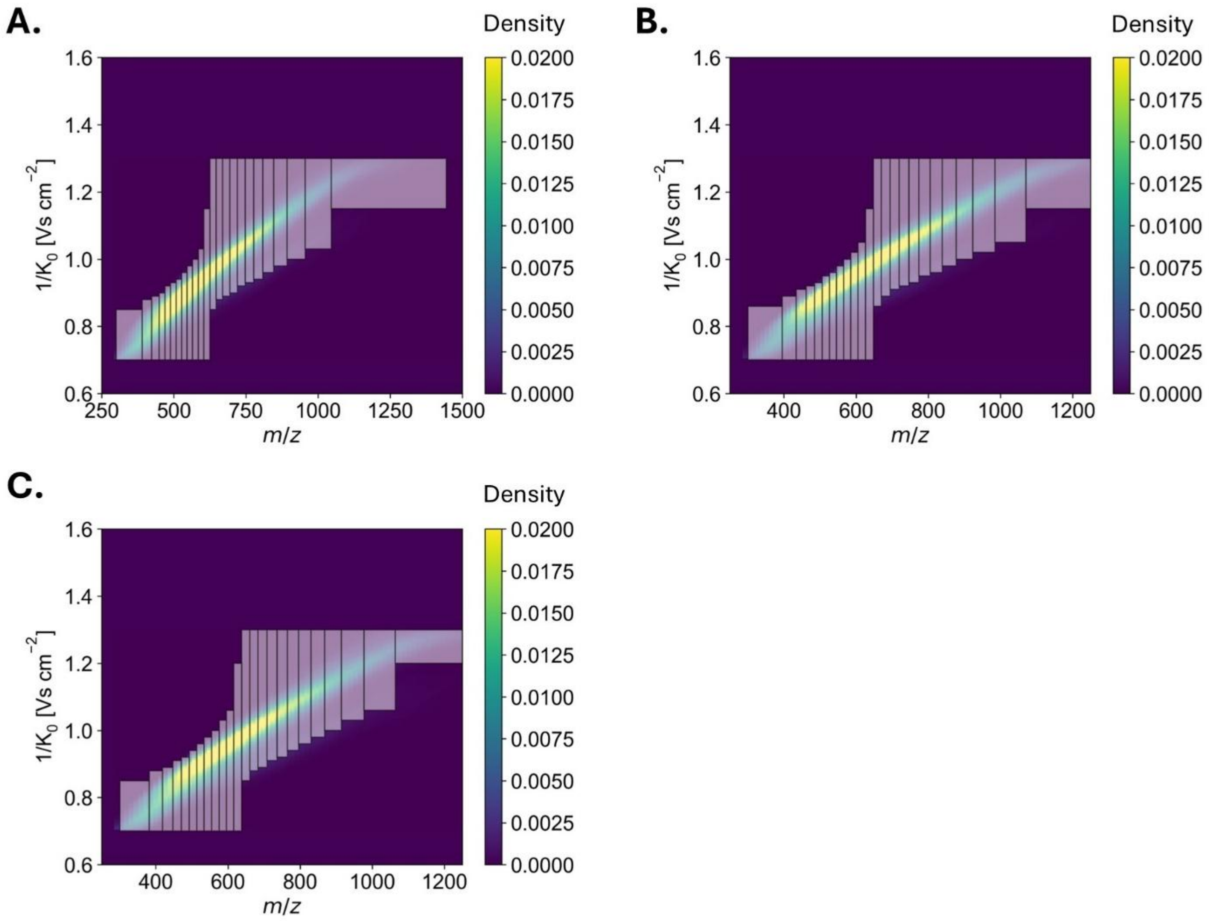


**Supplemental Figure S1**. **Optimal dia-PASEF acquisition schemes for dimethyl labeling based mDIA.** A 12-scan dia-PASEF method was used for acquiring data from dimethyl-labeled peptides in (A) the benchmarking experiment (EVs), (B) the application experiment (EVs), and (C) the application experiment (cells). Each scheme consists of one MS1 scan followed by 12 dia-PASEF scans with variable m/z isolation widths and two ion mobility windows within a single cycle. The color scale represents the density of precursors across the m/z and $1/K_0$ space.

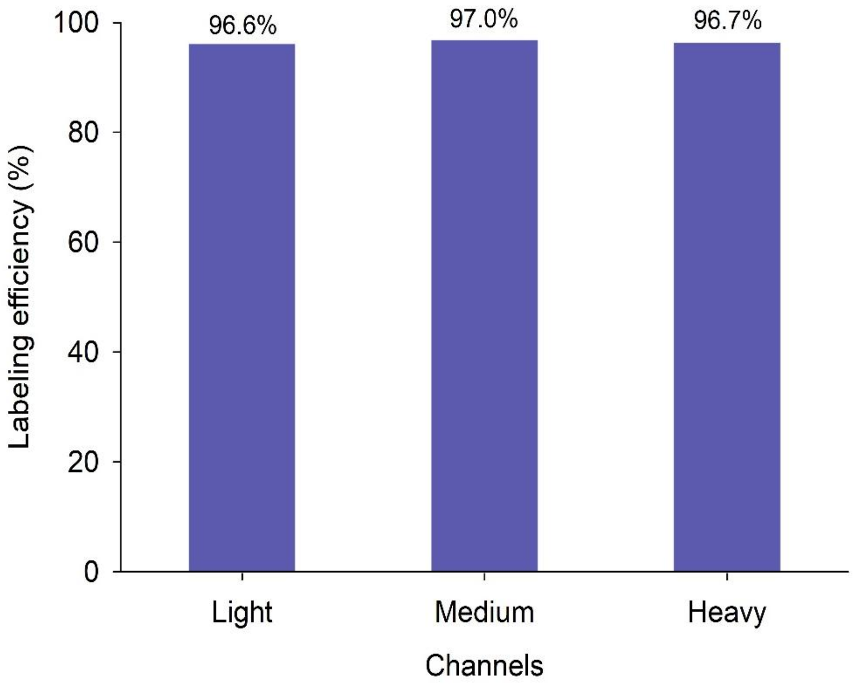


**Supplemental Figure S2. Dimethyl labeling efficiency**. Equal amounts of EV peptides were loaded into each dimethyl channel. The pooled sample was analyzed using DDA-PASEF and searched with FragPipe. Labeling efficiency was calculated based on the ratios of labeled peptides relative to all detected peptides (n = 1).

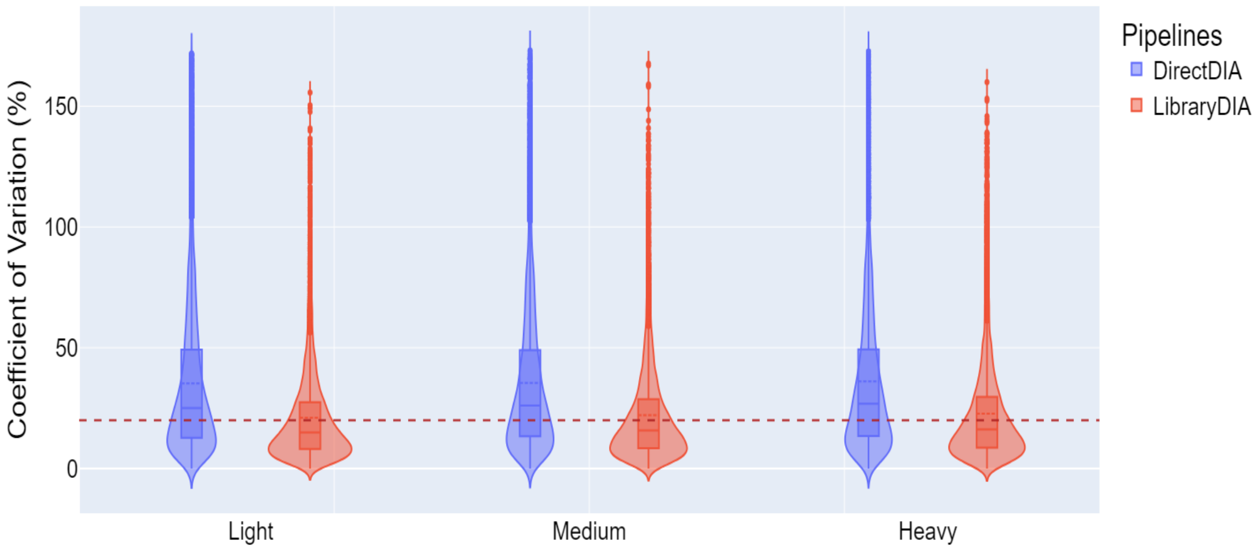


**Supplemental Figure S3**. **Comparison of quantification precision between library-based and library-free pipelines in the EV-based application experiment.** CV distributions for precursors quantified using the directDIA and library-based DIA pipelines in Spectronaut. Violin plots show the spread of CV%, with the red dashed line representing the 20% CV threshold.

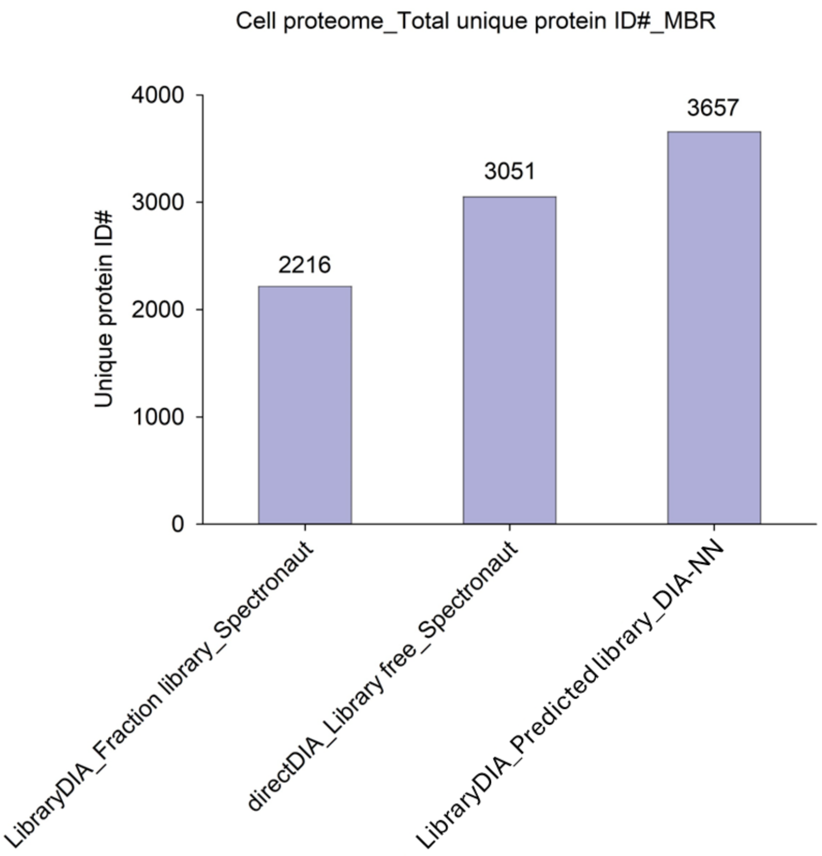


**Supplemental Figure S4**. **Protein identification numbers from the cell-based application experiment using different mDIA pipelines.** Each bar represents the total number of unique protein identifications across three channels in the match-between-runs (MBR) analysis (n=3). Fraction library is the project-specific library generated from fractions. Predicted library is from Thielert et al.

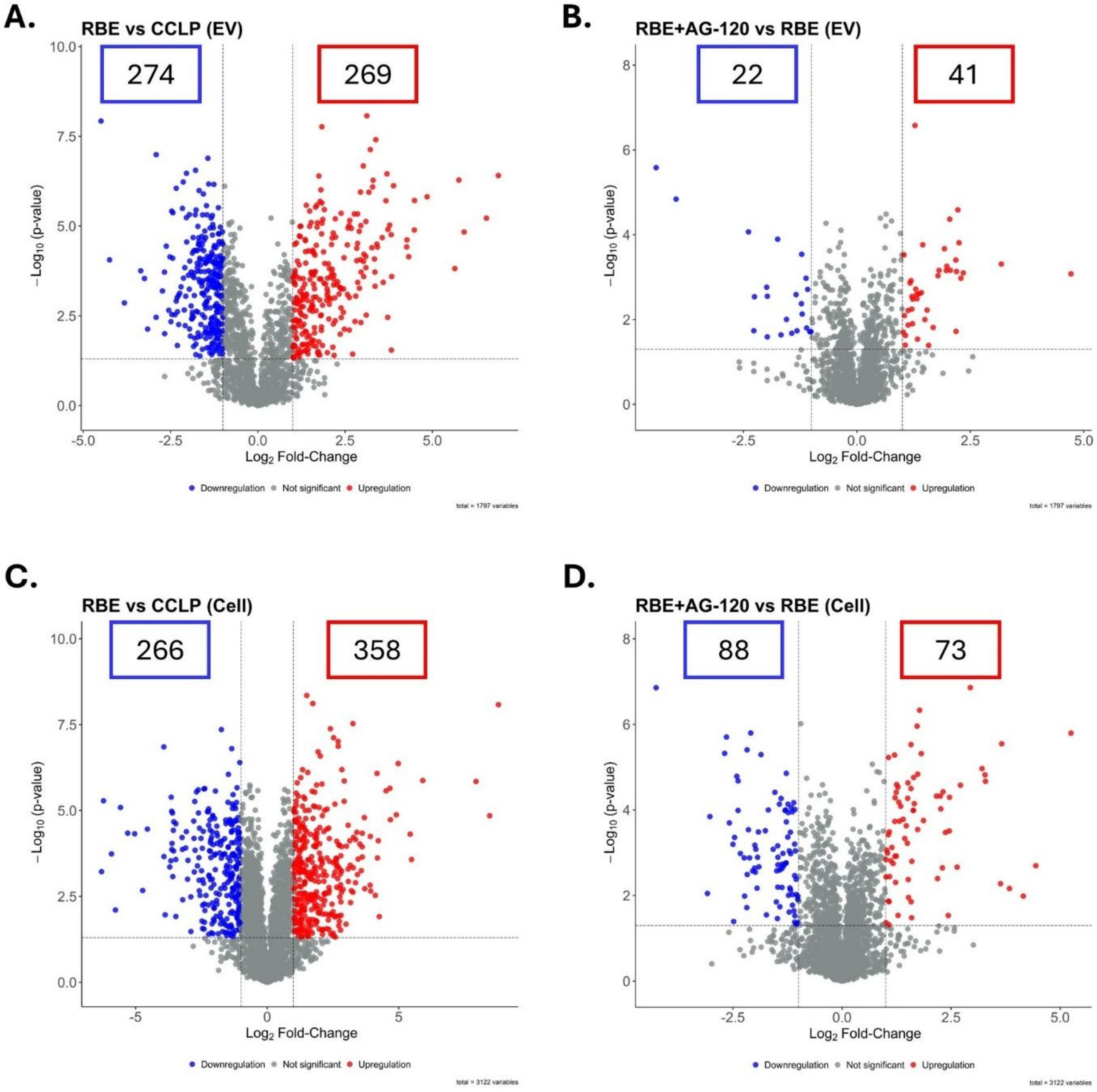


**Supplemental Figure S5. Significantly regulated proteins in cell- and EV-based experiments.** Volcano plots show comparisons of (A) RBE vs. CCLP in EVs, (B) RBE + AG-120 vs. RBE in EVs, (C) RBE vs. CCLP in cells, and (D) RBE + AG-120 vs. RBE in cells. Significance was determined using a two-sample Student's t-test with a p-value cutoff of 0.05 and an absolute $\log_2$ fold-change cutoff of 1. The numbers of significantly up- and down-regulated proteins are highlighted.

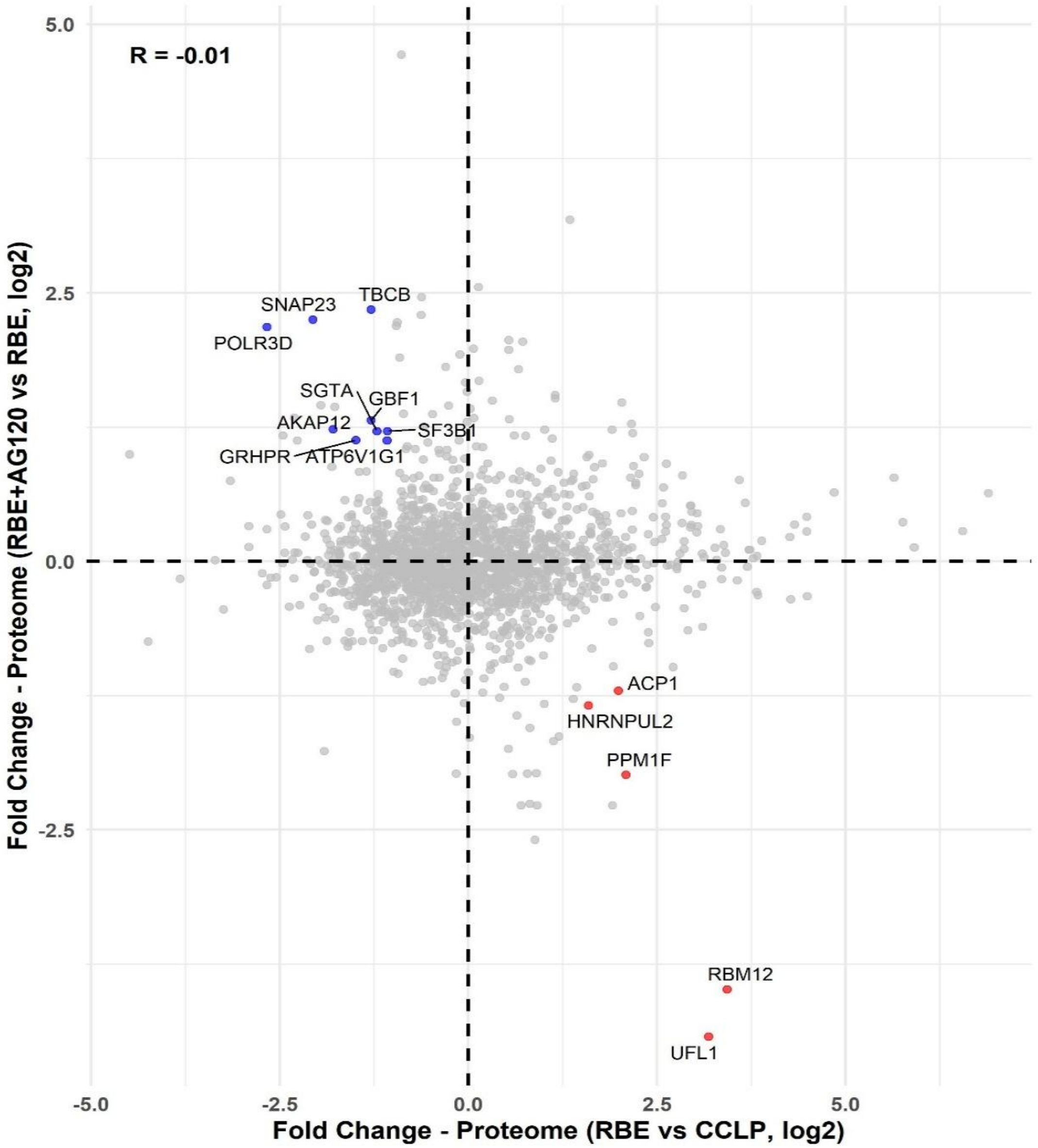


**Supplemental Figure S6. EV protein candidates for identifying IDH1 mutant iCCA subtype and monitoring AG-120 therapeutic response.** Scatter plot shows EV proteins with opposite expression patterns between IDH1 mutation and AG-120 treatment conditions. Proteins significantly upregulated in RBE vs. CCLP and significantly downregulated in RBE + AG-120 vs. RBE are shown in red, while proteins with the opposite pattern are shown in blue. Significance was determined using a two-sample Student's t-test with a p-value cutoff of 0.05 and an absolute $log_2$ fold-change cutoff of 1. All significant proteins are labeled.
